# Herpes zoster in HIV-1 infection: the role of CSF pleocytosis in secondary CSF escape and discordance

**DOI:** 10.1101/2020.01.28.922690

**Authors:** Lars Hagberg, Richard W. Price, Henrik Zetterberg, Dietmar Fuchs, Magnus Gisslén

## Abstract

HIV cerebrospinal fluid (CSF) escape is defined by a concentration of HIV-1 RNA in CSF above the lower limit of quantification of the employed assay and equal to or greater than the plasma HIV-1 RNA level in the presence of treatment-related plasma viral suppression, while CSF discordance is similarly defined by equal or higher CSF than plasma HIV-1 RNA in untreated individuals. During secondary CSF escape or discordance disproportionate CSF HIV-1 RNA develops in relation to another infection in addition to HIV-1. We performed a retrospective review of people living with HIV receiving clinical care at Sahlgrenska Infectious Diseases Clinic in Gothenburg, Sweden who developed uncomplicated herpes zoster (HZ) and underwent a research lumbar puncture (LP) within the ensuing 150 days. Based on treatment status and the relationship between CSF and plasma HIV-1 RNA concentrations, they were divided into 4 groups: *i)* antiretroviral treated with CSF escape (N=4), ii) treated without CSF escape (N=5), *iii)* untreated with CSF discordance (N=8), and *iv)* untreated without CSF discordance (N=8). We augmented these with two additional cases of secondary CSF escape related to neuroborreliosis and HSV-2 encephalitis and analyzed these two non-HZ cases for factors contributing to CSF HIV-1 RNA concentrations. HIV-1 CSF escape and discordance were associated with higher CSF white blood cell (WBC) counts than their non-escape (P=0.0087) and non-discordant (P=0.0017) counterparts, and the CSF WBC counts correlated with the CSF HIV-1 RNA levels in both the treated (P=0.0047) and untreated (P=0.002) group pairs. Moreover, the CSF WBC counts correlated with the CSF:plasma HIV-1 RNA ratios of the entire group of 27 subjects (P=<0.0001) indicating a strong effect of the CSF WBC count on the relation of the CSF to plasma HIV-1 RNA concentrations across the entire sample set. The inflammatory response to HZ and its augmenting effect on CSF HIV-1 RNA was found up to 5 months after the HZ outbreak in the cross-sectional sample and, was present for one year after HZ in one individual followed longitudinally. We find that HZ provides a ‘model’ of secondary CSF escape and discordance. Likely, the inflammatory response to HZ pathology within the CNS provoked local HIV-1 production by enhanced trafficking or activation of HIV-1-infected CD4+ T lymphocytes. Whereas treatment and other systemic factors determined the plasma HIV-1 RNA concentrations, in this setting the CSF WBC counts established the relation of the CSF HIV-1 RNA levels to this plasma set-point.

**Author summary:** Herpes zoster is a neurotropic infection common in people living with HIV. We studied if herpes zoster caused alterations in the cerebrospinal fluid viral (CSF) load when compared with blood levels in patients with and without antiretroviral treatment by studies of the cerebrospinal fluid. HIV-1 CSF escape and discordance were associated with higher CSF white blood cell counts than their non-escape (P=0.0087) and non-discordant (P=0.0017) counterparts, and the CSF white blood cell counts correlated with the CSF HIV-1 RNA levels in both the treated (P=0.0047) and untreated (P=0.002) group pairs. We found that the inflammatory response to herpes zoster pathology within the CSF provoked local HIV-1 production by enhanced trafficking or activation of HIV-1-infected CD4+ T lymphocytes. We suggest that herpes zoster provides a ‘model’ of secondary HIV CSF escape and HIV CSF/blood discordance, which increase the understanding of HIV central nervous system infection and might be of importance for eradication trials in sanctuary sites such as the central nervous system.

## Introduction

HIV-1 RNA is detected in the cerebrospinal fluid (CSF) throughout the course of untreated systemic infection, beginning early after initial exposure [1–3] and continuing until suppressed by antiretroviral therapy (ART) [4–7]. Among untreated people living with HIV infection (PLWH), only elite controllers consistently exhibit CSF HIV-1 RNA below the clinical laboratory lower limits of quantitation (LLoQ) [8, 9]. Though individual CSF:plasma HIV-1 relationships are variable, characteristically the CSF HIV-1 RNA concentrations in untreated PLWH average more than 10-fold lower than those of plasma, [5, 10, 11]. Combination ART is generally very effective in suppressing CNS HIV-1 infection, so that treatment that reduces plasma HIV-1 to below the clinical LLoQ is similarly effective in eliminating ‘clinically detectable’ CSF HIV-1 RNA. In our earlier studies of individuals ‘failing’ therapy, ART appeared to disproportionately lower CSF HIV-1 RNA compared to plasma[12, 13]. Even when more sensitive assays have been used to measure HIV-1 RNA, detection in the CSF was less common and at lower levels than that in plasma, consistent with the general treatment effectiveness on CNS infection[9, 14].

However, there are notable exceptions to this effective and proportionate effect of ART on CSF HIV-1 infection that have been collectively designated CSF escape in which the concentration of HIV-1 in CSF remains above the LLoQ and equal to or greater than that in plasma in the face of therapeutic plasma virus suppression [15–17]. This was coherently defined in a report of 11 treated patients presenting with a variety of neurological symptoms and signs who exhibited higher CSF than plasma HIV-1 RNA concentrations [18]. Additional reports preceded [19] and have followed [20–29], and this condition is now termed neurosymptomatic CSF escape based on the clinical presentation of neurological deficits in the presence of detectable CSF HIV-1 RNA despite systemic viral suppression[15, 17, 30]. This is clinically the most important type of CSF escape and can present with major neurological deficits and imaging abnormalities [20]. It at least partially overlaps with the neuropathological entity, CD8 encephalitis [31, 32], and involves replication of HIV-1 within the CNS despite systemic viral control. Unlike HIV-1 encephalitis that may complicate late untreated HIV-1 infection [33, 34], neurosymptomatic escape usually develops in individuals with blood CD4+ T lymphocyte counts above the 200 cells per µL threshold defining AIDS, and CD8+ T cell infiltration may be very prominent [31]. In these cases, it is presumed that the neurological symptoms and signs are causally related to CNS HIV-1 infection that is not effectively controlled by ART despite systemic efficacy, often in concert with a robust inflammatory reaction. This is supported by histopathological identification of HIV-1 in brain tissue of some cases, including some of those designated as CD8 encephalitis that share the CSF escape profile [32]. HIV-1 causality is also supported by the frequent therapeutic benefit of treatment modification on neurological symptoms and signs [18, 20, 29]. Most, but not all, of these individuals exhibit inadequate CNS treatment regimens related to drug resistance of the CNS virus population or to insufficient ‘penetration’ of the full combination of drugs into the CNS compartment by virtue of their pharmacological properties—or, frequently, to a combination of these two factors [18, 20, 27]. It may also involve important immunopathology with some features in common with the immune reconstitution inflammatory syndrome (IRIS) [32].

A second type of CSF escape, designated asymptomatic CSF escape, has also been documented in which CSF HIV-1 RNA is increased despite plasma suppression, but without clinical evidence of neurological injury. Asymptomatic CSF escape is more common than the neurosymptomatic form and seemingly largely benign [35]. Its reported incidence varies with the study population, the PCR assay used and its LLoQ, but generally has ranged near 5-10% of treatment-suppressed individuals [36, 37]. In most of these the escape appears to be transient or episodic and without evident clinical consequences, though CSF neopterin concentrations may be above those found in the presence of full CSF HIV-1 RNA suppression, indicating an inflammatory component associated with the asymptomatic escape[35, 36]. Unusual instances of more sustained asymptomatic escape have also been reported [36, 38]. Since these individuals do not have clear evidence of active neurological disease, asymptomatic escape is most commonly encountered in cohort studies in which lumbar punctures (LPs) and CSF analysis are components of the protocol, though also at times it can be an ‘incidental finding’ of a diagnostic LP.

The current report deals with a third type of CSF HIV-1 escape, termed secondary CSF escape, in which another nervous system infection appears to provoke, or at least associate with, elevation of the CSF HIV-1 RNA concentration despite plasma viral suppression, presumably as a consequence of the host local inflammatory responses to the inciting non-HIV-1 pathogen[15].

While more common in those with depressed blood CD4+ counts [39], herpes zoster (HZ) also presents in untreated PLWH at higher blood CD4+ T-cell counts than the more severe opportunistic infections defining an AIDS diagnosis [40]. Its incidence also appears to be higher in treated PLWH despite suppressive ART than in the non-HIV population [39, 40]. Beginning with reactivation in the sensory ganglion, varicella-zoster virus (VZV) spreads along the associated peripheral nerve both centrifugally to the associated dermatome, resulting in the characteristic rash, and centrally to the nerve root, and variably into the root entry zone of the spinal cord or brain stem [41]. This is accompanied by reactive inflammation that may extend from the ganglioneuritis to the adjacent meninges, with common but variable CSF pleocytosis [42]. Fortunately, HZ is usually self-limiting in both untreated and treated HIV-1 infection, though not always, and more severe systemic dissemination and nervous system spread have been reported in immunosuppressed patients [43–47]. Because early antiviral treatment may shorten the eruption and reduce the incidence of postherpetic neuralgia and other complications, it is recommended in current NIH Guidelines (https://aidsinfo.nih.gov/guidelines/html/4/adult-and-adolescent-opportunistic-infection/392/whats-new-in-the-guidelines).

Having noted some cases of secondary escape in individuals with HZ in our HIV-1 cohort study and in the course of HIV-1 clinical care in Gothenburg, we undertook a review of all the cases of HZ in HIV-1-infected individuals who underwent lumbar puncture (LP) in the longitudinal Gothenburg HIV CSF Study [48, 49] within 5 months of the onset of the HZ rash. To date, 734 PLHW have been included in this study. The protocol includes LP at diagnosis, 3 months after treatment initiation and thereafter yearly. Extra samples have been obtained in settings of opportunistic infections and HZ. It was thus possible to retrospectively include individuals on ART with and without CSF escape, and to extend the analysis to untreated individuals in whom CSF HIV-1 RNA concentrations exceeded those of plasma (defined here as discordant CSF:plasma ratio) and those with the more common finding of lower CSF than plasma HIV-1 concentrations (nondiscordant CSF HIV-1 RNA). We reviewed this experience to identify factors that might predispose to higher HIV-1 concentrations in CSF than plasma in the presence and absence of ART, *i.e.*, in individuals with CSF escape and CSF discordance.

The prominent inflammatory aspect of HZ provides a relatively common setting to examine the effects of focal inflammation within the neuraxis on CSF HIV-1 concentrations, and we therefore undertook a review of our experience with HZ in PLWH who underwent LP and CSF analysis.

## Methods

### Design

This was a retrospective study drawn from the records of the Gothenburg Infectious Diseases Clinic, searched for HIV-1 infection, HZ rash and LP. All records were reviewed by one of us (LH), and those with an LP within 150 days of the onset of the dermatomal rash were included. All LPs were performed within the Gothenburg HIV CSF Study Cohort protocol, approved by the institutional IRB and with informed consent. The identified individuals with HZ were classified by treatment status and the relation of CSF to plasma HIV-1 RNA concentrations into four groups using the following definitions:

#### On ART

***Secondary CSF escape***: a) on chronic stable and consistent ART treatment (at least 8 months of therapy); b) plasma HIV RNA <500 copies/mL; c) CSF HIV-1 RNA concentrations above the clinical laboratory LLoQ of 20 copies/mL; and d) CSF ≥ plasma HIV-1 RNA.

***On therapy no escape***: as in a) above; but b) CSF HIV-1 RNA below the laboratory LLoQ, or CSF < plasma HIV-1 RNA.

#### Off ART

***CSF discordance***: a) untreated (naïve or off ART >8 months); and b) CSF ≥ plasma HIV-1 RNA.

***No CSF discordance***: untreated as in a) above; but b) with CSF < plasma HIV-1 RNA.

Cross-sectional analysis included data from the first LP after the onset of HZ. CSF and plasma HIV-1 RNA were measured in all subjects using the Roche Cobas Taqman assay with the LLoQ of 20 copies/mL; a value of 19 copies/mL was assigned as the default value for any levels below 20 in calculations and descriptive statistics. The CSF:plasma ratio was calculated using log_10_ values: *log_10_ CSF HIV-1 RNA – log_10_ plasma HIV-1 RNA*.

ART regimen at the times of HZ outbreak and CSF sampling were available for all subjects. CSF white blood cell (WBC) counts, concentrations of CSF and blood neopterin and CSF neurofilament light chain protein (NfL) were measured for most using previously reported methods [50, 51]. The time from rash outbreak to LP was recorded along with notation of the location of the dermatomal rash. A simple scale was used to compare the distance of the rash from the lumbosacral LP: *1* for lumbosacral dermatome, *2* for thoracic, *3* for cervical and *4* for cranial nerve. Selected cases in which CSF HIV-1 measurements were available before and after the HZ episode were reviewed for longitudinal findings.

To supplement the experience with secondary escape, in selected analyses we included two additional individuals with secondary CSF escape related to other infections: one with neuroborreliosis (common in the Gothenburg area and documented by intrathecal antibody production) and an uncommon case of HSV-2 encephalitis documented by CSF PCR.

### Statistics

Descriptive statistics, comparisons of subgroups and analysis of correlations among variables generally used nonparametric methods, with linear regressions performed in selected analyses. These along with graphic presentations were all performed using Prism 8.3 (GraphPad Software). For some analyses, data were log_10_ transformed; in the case of CSF WBCs, when the initial laboratory value was zero, we substituted a value of one for analysis and graphing of the transformed data.

## Results

### Cross-sectional study population

We identified 25 HIV-1-infected individuals who underwent LP within 150 days after the onset of HZ rash: 9 on and 16 off ART (Table 1). These were divided into the 4 categories defined above by treatment status and the relation of CSF to plasma HIV-1 RNA levels. Most were treated with a course of valacyclovir or acyclovir after recognition of the rash. Table 1 lists the salient background data for each of the subjects grouped according to these four categories. The neurological symptoms and signs in the HZ patients were consistent with the zoster itself, and none showed overt evidence of extension of the infection into the CNS or more remote dissemination. All recovered without neurological sequelae beyond sensory disturbances. The two additional treated individuals with secondary CSF escape related to other infections (neuroborreliosis and HSV-2) are included at the bottom of the Table 1. In those with HZ the median age was 42 years and the median time from rash outbreak to LP was 18 days with a range from one to 150 days. The patient with neuroborreliosis was a bit older, at 62 years. In general, the demographics, including sex, race and ethnicity, reflected those of HIV-1 infection in Sweden and of the Gothenburg cohort. There were 16 men and 9 women including 18 Caucasian, 6 Black-African, 2 Asian and one Latin-Hispanic.

### HIV-1 RNA concentrations in the four groups

Figure 1 depicts the relations between CSF and plasma HIV-1 RNA concentrations for each individual within their classified group, with the two non-HZ patients included among those with HZ CSF secondary escape (Figure 1A). The lines connecting the plasma to CSF HIV-1 RNA concentrations of each individual emphasize the relationships of the HIV-1 RNA in the two fluids, higher in the CSF than plasma in the escape and discordant groups (left panels, A and C) and lower in the no escape and nondiscordant groups (right panels, B and D). The nondiscordant group exhibited the more typical negative CSF:plasma ratio, here with a more than 10-fold mean higher plasma than CSF HIV-1 RNA concentration. In accordance with the grouping definition, the plasma HIV-1 RNAs in the treated groups were similar. Only one of the escape groups was not suppressed to the laboratory LLoQ of ≤20 HIV-1 RNA copies per mL (or 1.30 log10 copies per mL) in plasma, but the mean CSF HIV-1 RNA in the HZ escape group was 2.68 log_10_ copies per mL, a modest but distinct level of CSF escape from therapy, particularly in comparison to the fully suppressed CSF levels in the no escape group. As expected, the plasma HIV-1 RNA concentrations were higher in the two untreated groups (mean of 4.69 log_10_ copies per mL in the discordant and 5.14 log_10_ copies per mL in nondiscordant) but with the CSF distinctly higher than its companion plasma in the discordant samples while the plasma HIV-1 RNA was higher than that of the CSF in the nondiscordant group. This figure not only shows the relationships between CSF and plasma HIV-1 RNA that conform to their group definitions but emphasizes the distinct differences among these four groups that serves as the basis for comparing their other characteristics.

**Figure 1.**
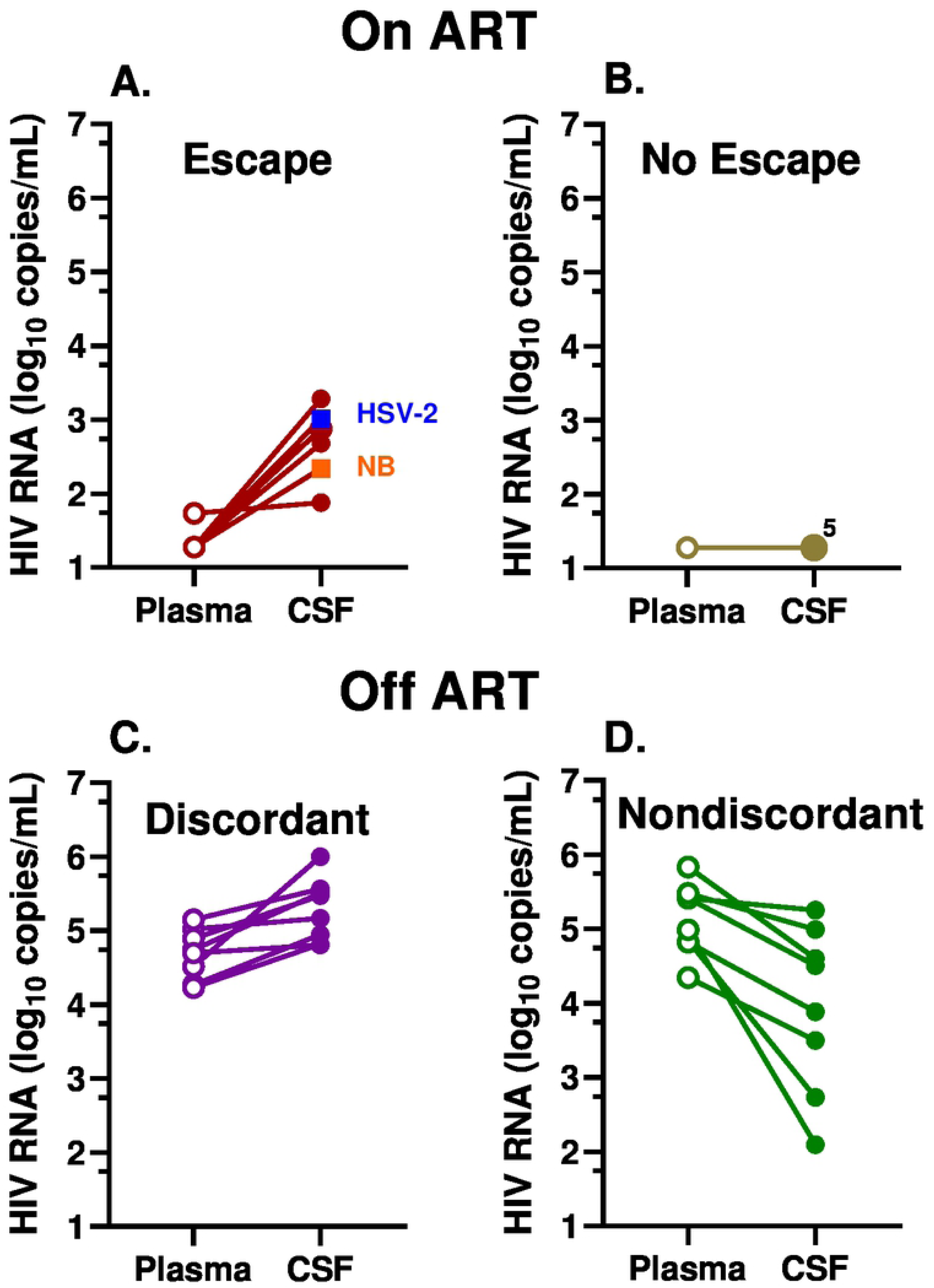
Relation between plasma and CSF HIV-1 RNA concentrations in the 4 subject subgroups. **A.** Treated CSF escape group including the 4 HZ (brown circles) and two other infections—neuroborreliosis, *NB* (orange) and HSV-2 encephalitis (blue); **B**. Treated, no escape, 5 HZ with overlapping CSF HIV-1 RNA levels at the LLoQ (pale green); **C.** Untreated CSF discordant (purple); and **D.** Untreated CSF nondiscordant group (dark green). Plasma values are represented by open symbols and CSF with solid symbols and lines connect plasma and CSF from the same individual.

In order to better understand the reversal of CSF:plasma HIV-1 RNA ratios by HZ in some, but not all, of the identified individuals, we examined a number of other variables, including the CSF WBC count, CSF neopterin concentration, blood CD4+ T-lymphocyte counts, CSF NfL concentration, time between the HZ rash outbreak and CSF sampling, and distance between the HZ outbreak and the site of LP (Figure 2).

**Figure 2.**
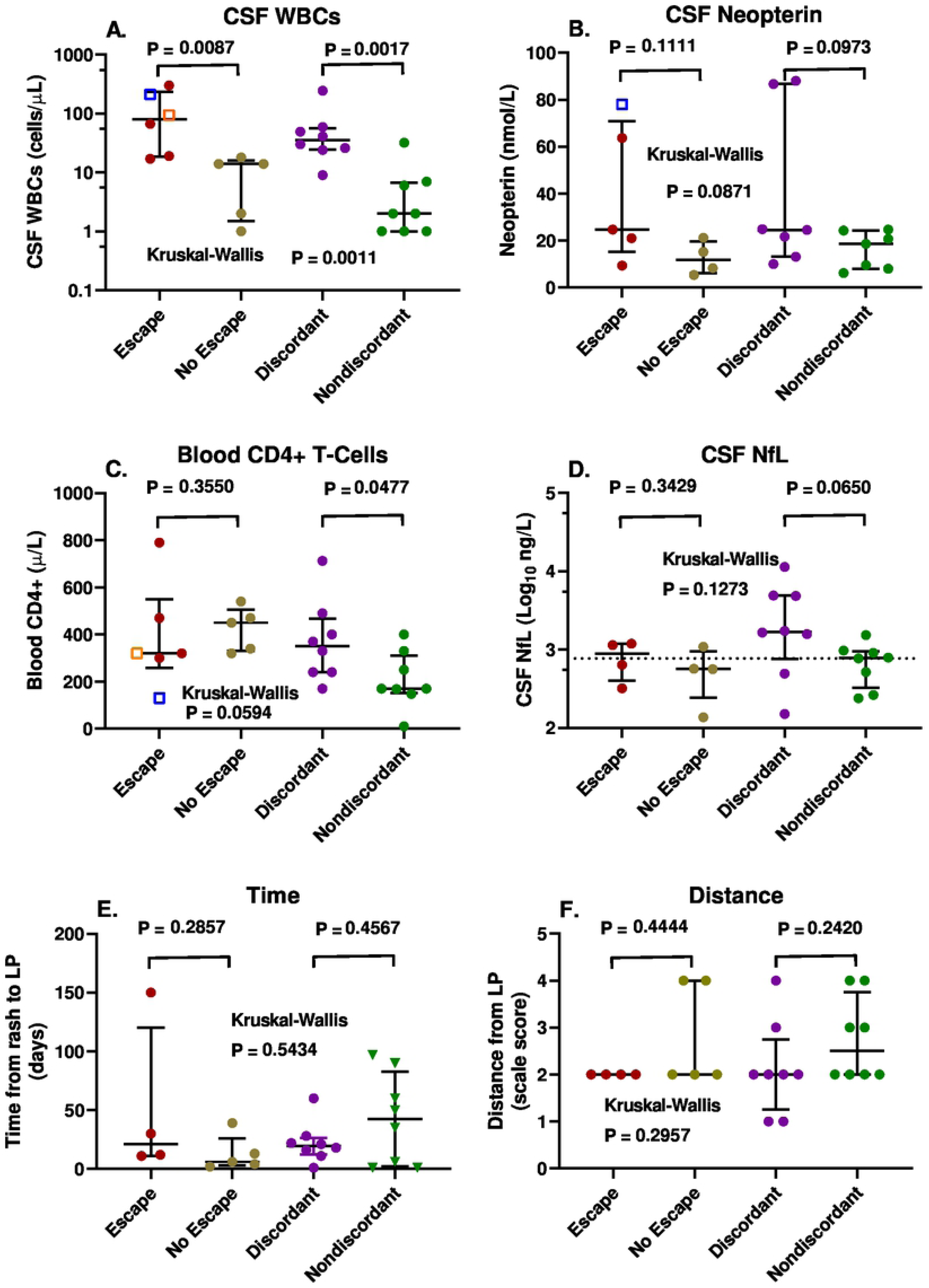
Comparisons of six variables among the four subject groups. Symbol colors are the same as in Figure 1 above. **A.** CSF WBC counts with escape group including the 2 non-HZ infections; **B.** CSF neopterin concentrations with escape group including the HSV-2 encephalitis; **C.** blood CD4+ T cell counts with escape group including the 2 non-HZ infections; **D.** CSF NfL concentrations in HZ; **E.** Time from outbreak of HZ rash to LP; and **F.** scaled distance from HZ eruption to LP site. Error bars show medians and interquartile ranges. The statistical comparisons shown are between the treated (escape and no escape) and untreated (discordant and nondiscordant) group pairs by Mann-Whitney test.

### CSF WBC count

The CSF WBC counts differed in the escape (including the two non-HZ cases) and discordant groups with CSF ≥ plasma HIV-1 RNA concentrations compared to their non-escape and nondiscordant counterparts (Figure 2A), so that higher CSF WBC counts were associated with higher CSF HIV-1 than plasma HIV-1 RNA levels in both of these group pairings.

In order to examine this further, we separately assessed the relationship of the CSF HIV-1 RNA levels to the CSF WBC counts in the treated (escape and no escape) and untreated (discordant and non-discordant) groups using linear regression after log_10_ transformations (Figure 3A). This showed a strong correlation within each of these two paired groups, indicating an effect on the CSF HIV-1 RNA concentrations across the range of CSF WBC counts in both the treated and untreated individuals. Both the escape and discordant individuals were in the higher CSF WBC range while the non-escape and non-discordant were in the lower CSF WBC range of each of these lines which were separated by nearly 3 log_10_ copies/mL of HIV-1 RNA related to the effect of systemic treatment.

**Figure 3.**
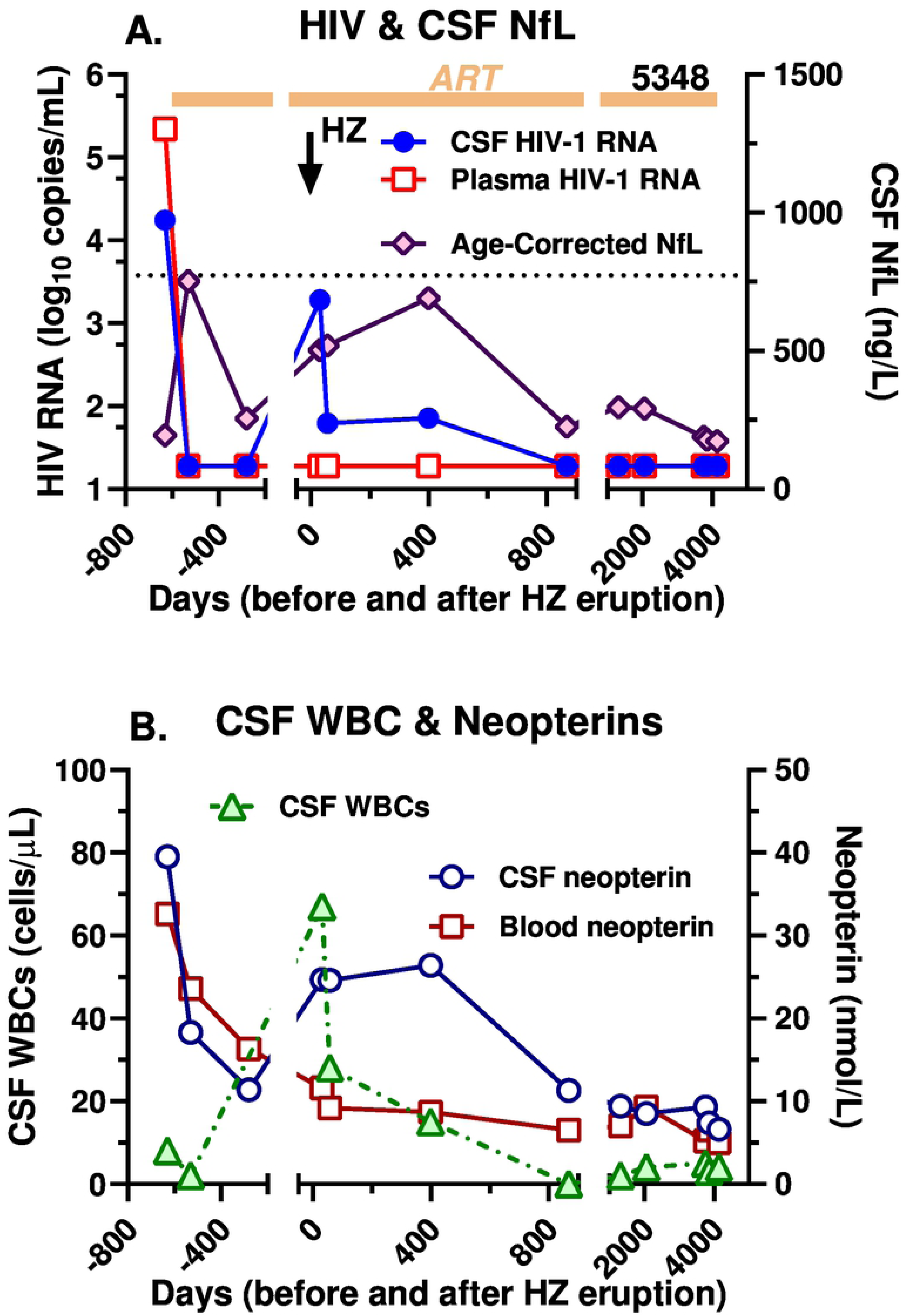
Relation of CSF HIV-1 RNA and CSF:plasma HIV-RNA ratio to CSF WBC counts. **A.** Linear regressions plotting the relation of the log_10_ CSF HIV-1 RNA concentrations per mL to the log_10_ CSF WBC counts per µL in the treated (lower line) and untreated (upper line) subjects. Symbol colors are as in Figure 1 as per legend. The larger symbols for the no escape subjects indicate two points with overlapping values. **B.** Linear regression plotting the relation of the CSF:plasma HIV-1 RNA ratio (calculated as log_10_ difference between CSF and plasma HIV-1 RNA values) to the log_10_ CSF WBC counts per µL. The horizontal dotted line indicates equivalent values in CSF and plasma. Open symbols are the same colors as in Figure 1. Statistics shown are the P and R squared values from the linear regressions which are plotted with 95% confidence intervals.

Because of this separation between the two sample pairs, we next examined the correlation of the CSF WBC counts with the ratios of CSF to plasma HIV-1 RNA (expressed as the log_10_ difference between CSF and plasma HIV-1 RNA) across the entire sample set as shown in Figure 3B. This revealed a strong, correlation that included all four subject groups along a single regression slope. Taken together these data suggest that the plasma HIV-1 (whether treated or untreated) set the overall level of infection in the plasma, but that the CSF WBC response then strongly influenced the relationship of the CSF HIV-1 RNA concentration to that plasma level. This WBC effect resulted in a ‘reversed’ CSF-plasma ratio (*i.e*., ≥1) at a mean CSF WBC count of about 16 cells per µL where the regression crossed the line of no difference between CSF and plasma HIV-1 RNA depicted by the horizontal dotted line in Figure 3B.

### CSF neopterin

The Gothenburg cohort study included measurement of CSF neopterin at most LP visits. THis is a sensitive measure of enhanced immune activation-inflammation in the CNS during HIV-1 infection [50]. In this small series, the CSF neopterin concentrations were above normal in most of those with HZ and higher in the escape and discordant groups than their non-escape and non-discordant counterparts, mainly related to two high values in each of these groups, and overall the group pairs were not significantly different (Figure 2B). The CSF neopterin did correlate with the CSF:plasma HIV-1 ratio, though not as strongly as the CSF WBC count (linear regression P = 0.0184; R^2^ = 0.231). Thus, CSF neopterin measurements supported the association of inflammation with the CSF relation to plasma HIV-1 RNA, but were not as clear a predictor as the CSF WBC count, though the CSF neopterin and WBC counts, themselves, were strongly correlated (P= 0.0001; Spearman r =0.7195).

### Blood CD4+ T-lymphocyte count

The blood CD4+ T-cell counts of the HZ patients were mostly above the levels predisposing to opportunistic infections, with a median near 400 cells/µL in the treated groups and 350 cells/µL in the discordant group off ART (Figure 2C). They were lower in the nondiscordant group off therapy with a median near 200 cells/ µL. Only in this nondiscordant did the CSF HIV-1 RNA correlate with the CD4 count (P = 0.0024, Spearman). Mechanistically, this may have related to an impact of low CD4 levels on the CSF WBC counts, since in the untreated the CSF WBC counts correlated with the CD4 counts (P=0.004) in contrast to the absence of significant correlation in the treated subjects (P=0.081). This is perhaps consistent with previous observations that lower CSF HIV-1 RNA levels and WBC counts are found in individuals with CD4 counts <50 cells/µL [5] but suggests that this effect may extend to blood CD4+ T-cell counts above 50 cells/µL.

### CSF NfL, time and distance from rash outbreak

The Gothenburg cohort study also measured CSF NfL at most visits. Because HZ can elevate the CSF NfL presumably as a result of axonal injury of nerve roots and perhaps at times spinal cord [52], we reasoned that the CSF NfL levels in this setting would reflect the activity and severity of the HZ ganglioneuritis and perhaps its distance from the LP sampling site. We also more directly examined the timing of the LP in relation to the rash outbreak and the distance of the rash from the LP sampling site using the simple scale described above to see if any of these measures correlated with CSF HIV-1 RNA. However, none showed a clear correlation (Figure 2D-F). While the severity and timing of the ganglioneuritis and its distance from the LP site are likely interactive and interrelated, with the limited sampling of this cross-sectional study we did not show a clear individual relation to the CSF HIV-1 RNA or its relation to plasma HIV-1 RNA. If these factors did indeed exert an effect on the CSF HIV-1 RNA, it was not as robust as the CSF WBC reaction.

#### Longitudinal observation

CSF and blood samples from one of the discordant subjects with HZ (SID #5348 in the Table) were obtained at several study visits before and after the index HZ LP included in the cross-sectional analysis (Figure 4). He developed HZ in February 2006, nearly two years after starting suppressive ART. At the time of the index LP 30 days after the onset of zoster rash, his CD4+ T-lymphocyte count had risen from a nadir of 40 to 300 cells/µL, his plasma HIV-1 RNA was suppressed to <20 copies/mL, but his CSF HIV-1 RNA was elevated at 1,920 copies/mL despite having been undetectable for at least a year and one-half. The CSF HIV-1 RNA then remained elevated for 400 days at 62 and 72 copies/mL but returned to below the detection limit by the next LP at 868 days and through the duration of follow-up out to 11 years later (Figure 4A). The plasma HIV-1 RNA remained below the LLoQ throughout this long period of observation. The CSF NfL concentration rose early after initial therapy, but then decreased to its basal level before it again rose during the HZ episode, remaining above the basal level through 400 days before returning to baseline after longer follow-up; none of these measurements were actually above the upper limit of age-corrected normal levels of NfL [53]. This presumably reflected the protracted course of zoster ganglioneuritis (Figure 4A). The CSF WBC count followed a time course very similar to that of the CSF HIV-1 RNA concentration, increasing initially to 67 cells/mL at the index HZ LP, and then subsiding to 28 and 15 cells/mL over the ensuing year before returning to normal along with the CSF HIV-1 RNA and NfL (Figure 4B). CSF neopterin was elevated above the plasma concentration before treatment, fell with ART but then also rose over the course of the HZ before falling again after 2 years. While the plasma neopterin was also elevated before treatment initiation and fell with ART, it did not change its trajectory appreciably during the HZ episode, supporting the compartmentalized character of the protracted HZ course, its associated inflammation and effect on local HIV-1 replication. In summary, the markers of inflammation (CSF WBCs and neopterin), axonal injury (CSF NFL) and local intrathecal HIV-1 infection (CSF HIV-1 RNA) all followed a similar temporal pattern during the HZ episode, linking these processes to the VZV reactivation and spread, and the inflammatory response over a prolonged time period.

**Figure 4.**
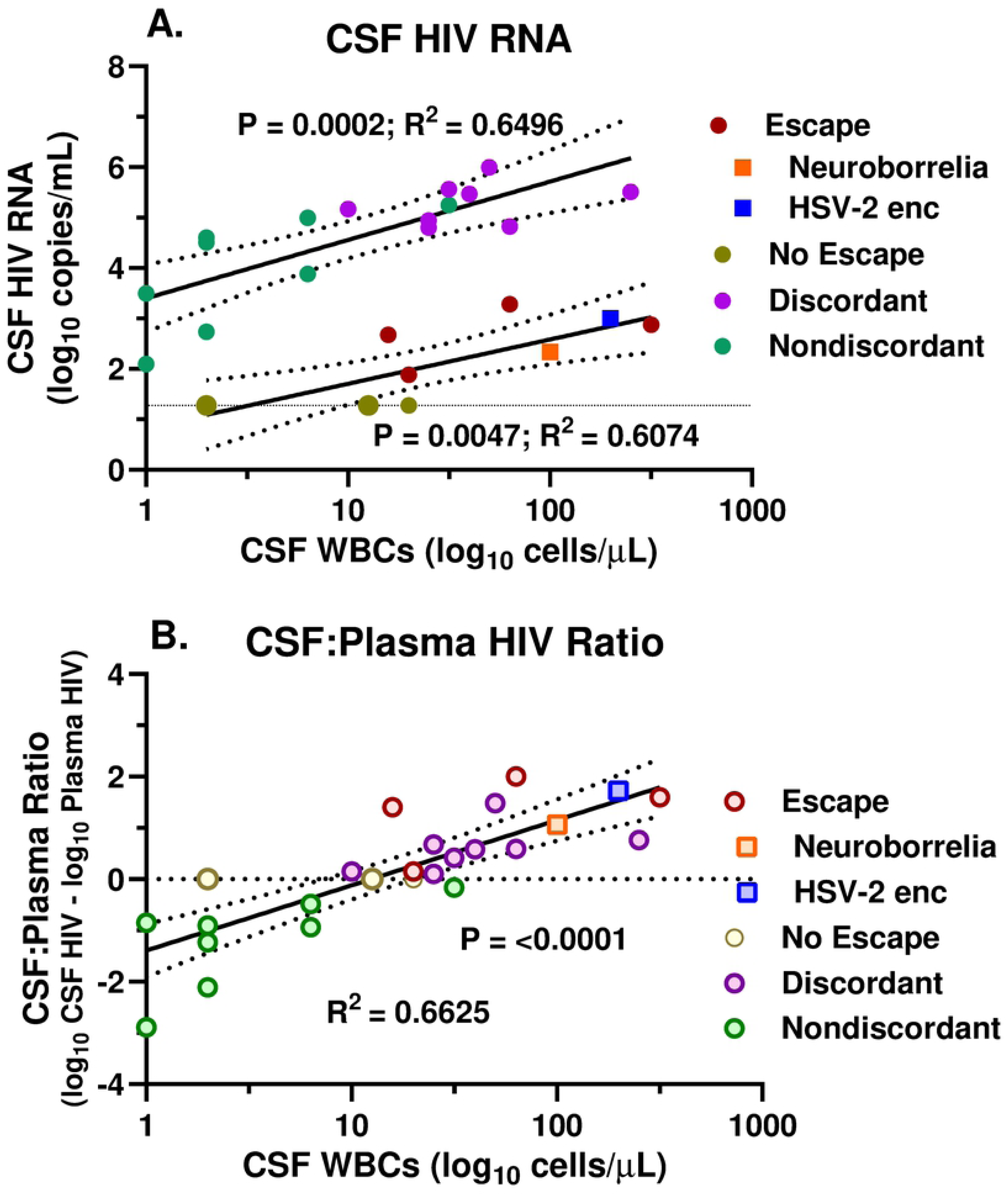
Longitudinal course in an individual with HZ and secondary CSF escape. The two panels plot the long-term longitudinal course in HZ CSF escape subject 5348 from before initial treatment, through the HZ episode, to more than a decade later. **A.** shows the course of CSF and plasma HIV-1 RNA concentrations and age-corrected NfL. Most notable is the rise in CSF HIV-1 RNA after the HZ (time 0 designated by the downward vertical arrow) and its continued elevation for more than one year despite continued plasma HIV-1 suppression; CSF NfL also rose above baseline levels during this period. **B.** CSF WBC and CSF and blood neopterin over this same time course. The CSF WBC count rose after the HZ and remained abnormal for over a year, similar to the CSF HIV-1 RNA in panel A. The CSF neopterin also increased during the HZ episode and remained relatively high over the ensuing year before falling later while the plasma neopterin did not alter its trajectory with the HZ episode.

## Discussion

This retrospective analysis indicates that HZ was associated with an increase in local HIV-1 production within the CSF compartment and strongly suggests that this was mediated through the local inflammatory response to VZV reactivation, and particularly by the lymphocytic pleocytosis. In this series of individuals, a more robust inflammatory response associated with an increase in CSF HIV-1 RNA concentration in relation to that of plasma HIV-1 RNA both in those on ART with secondary CSF escape and those off treatment with CSF discordance. Our analysis suggests that this augmentation of local HIV-1 production was associated with the WBC inflammatory cell response to HZ that then determined the level of CSF HIV-1 RNA in relation to that of plasma in both treatment and nontreatment settings. Our findings thus support the concept of secondary CSF escape, suggest an important mechanistic component, and have implications for other types of CSF escape and, more broadly, for one of the mechanisms that can contribute HIV-1 infection in CSF in other settings.

Both CSF escape and discordance are characterized by a ‘disproportionate’ CSF HIV-1 RNA concentration compared to that of plasma in the treatment and non-treatment settings, respectively. These relationships of CSF to plasma HIV-1 are unusual. In the absence of treatment CSF HIV-1 RNA levels are usually about 10-fold or more lower than those in plasma, while ART that effectively suppresses systemic viremia is usually similarly effective in suppressing CSF HIV-1 RNA [4, 13]; even in cases of systemic treatment failure plasma HIV-1 RNA may exceed that of CSF by more than the 10-fold ration [6]. CSF discordance in untreated and CSF escape in treated PLWH thus provide important examples of the reversal of the usual ratio of CSF to plasma HIV-1 concentrations. Indeed these relationships have been previously reported. CSF escape has been reported in neurologically complicated HZ [45] and discordance reported in a case of HZ meningitis[54] and a case of HZ myelopathy with compartmentalization of CSF HIV-1 sequences [55].

Our findings establish secondary escape as a third type of CSF escape joining the previously characterized neurosymptomatic and asymptomatic clinical forms of CSF escape discussed earlier [15-18, 20, 36]. It also suggests similar mechanisms underlying CSF escape and discordance in this setting. In contrast to the other two forms, secondary HIV escape has not been well characterized, and this report represents one of the first to examine a group of examples. In our 4 HZ subjects and the two without HZ (neuroborreliosis and HSV-2 encephalitis) with secondary escape, the CSF HIV-1 concentrations did not distinguish them from the broad range of either symptomatic or asymptomatic escape. In these individuals the CSF HIV-1 escape was also asymptomatic or at least obscured by the main presenting infection. There were no additional neurological symptoms and signs (for example, no headache or stiff neck to suggest meningitis, nor cognitive impairment to indicate HIV-1 encephalitis) that could not be attributed to the VZV eruption, or to the borrelia or HSV-2 CNS invasion, though in the latter case, this would have been difficult. These same considerations generally applied to the untreated individuals with HZ and discordant CSF HIV-1 RNA levels who also had no symptoms or signs beyond those attributable to HZ.

Our case series was small, but overall more than two-thirds of our PLWH and HZ exhibited pleocytosis (defined as a CSF WBC counts >5 cells/µL), a higher frequency than the nearly 50% reported in non-HIV patients [42]. In our subjects without escape or discordance the frequency of pleocytosis was similar to that reported in non-HIV subjects, 3 of 5 and 4 of 8 respectively. However, in the escape and discordance groups all had abnormal CSF WBC counts (median of 43 and 60) that were higher than in the non-escape and non-discordant groups (median of 14 and 4 cells/µL).

Common to both the treated and untreated groups, there was a relationship of the CSF WBC count to the CSF viral load and to the ratio of CSF to plasma HIV-1 RNA concentrations. Both treated and untreated groups showed a threshold between 10-30 cells/µL at which this ratio transitioned to higher CSF than plasma HIV-1 RNA concentrations that defined CSF escape or discordance depending on treatment status. The impact of the CSF WBC count on CSF HIV-1 infection in both treated and untreated individuals was most evident in relation to the CSF:plasma HIV-1 RNA ratio in the entire group of samples as depicted in Figure 3B. These data argue that while systemic factors and therapy established the overall level of infection and the plasma HIV-1 concentration, the CSF WBC count (comprised mainly of CD4+ and CD8+ T lymphocytes [56, 57]) strongly influenced the relationship of the CSF HIV-1 RNA concentration to the plasma level. This WBC effect resulted in a ‘reversed’ CSF-plasma ratio above a mean of about 16 WBCs per µL where the regression line in the figure crossed the line of no difference between CSF and plasma HIV-1 RNA (Figure 3B). This analysis also argues for the mechanistic similarity of CSF escape and discordance in this setting.

HZ is an inflammatory disorder in those without and with HIV-1 infection that begins in the dorsal root ganglion, spreads centrifugally along the peripheral nerve, and seeds the skin of the related dermatome, leading to the characteristic rash. Infection also spreads centripetally to the nerve root and, to a variable degree, beyond, into the spinal cord or brain stem [58–60]. It also may spill over into the leptomeninges, producing an ‘aseptic’ meningitis.

Our leading hypothesis in explaining the development of CSF escape and discordance in this setting is that HZ provokes local inflammation along this pathway that includes CD4+ T-cells, an important component in VZV defense [61–63]. Some of these CD4+ cells are most likely VZV-specific, but others may also include bystander cells with other antigen specificities responding to the heightened inflammatory milieu. It remains to be seen which population of these cells contains the HIV-1 provirus. With activation these cells may exhibit clonal expansion releasing HIV-1 of limited genotypes or they may initiate a broader local HIV-1 infection with other activated CD4+ T-cells providing targets for further local amplification [64–67]. In the presence of ART this replication, and particularly local amplification, might be constrained, though sufficient to produce virions detectable in the CSF. Indeed, the role of drug resistance may be important in understanding secondary escape. Evaluation of these individuals did not include CSF HIV-1 drug resistance testing. However, in the treated individuals undergoing LPs before or after the HZ episode, CSF HIV-1, like its plasma counterpart, was well controlled by the individual’s ART regimen. The output of virus in what had previously been and would again later be ‘adequate’ treatment may argue for clonal CD4+ expansion and HIV-1 production rather than ongoing local replication and amplification. While this needs to be directly tested, it might suggest that drug resistance was not as critical factor in secondary escape as it frequently is in neurosymptomatic CSF escape [18, 20]. In a case of complicated HZ with CSF discordance that provides precedence for those in our report, Falcone and colleagues found compartmentalization of CSF HIV-1 RNA and suggested that the CSF population included X4 lymphotropic phenotypes based on sequence analysis (though sequences were not listed in the publication) [55]. Further analysis of such cases should be informative on this issue.

Here, as in other settings, a crucial question is whether CSF pleocytosis was the cause of or the response to intrathecal HIV-1 infection. Following the arguments above, we favor the former: that the CSF WBCs importantly contributed *to* the level of CSF HIV-1 RNA. In the HZ setting (and the two other included infections) the secondary infection within the neuraxis was likely a sufficient explanation for the CSF pleocytosis, and the level of this CSF WBC response then led to the enhanced CSF HIV-1 RNA production. It is, however, theoretically possible that the local HIV-1 replication initiated by the HZ then further augmented the T-cell response, and that this included HIV-1-specific T-cells that carried the HIV-1 provirus and subsequently contributed further to the HIV-1 RNA burden in a self-limited positive feedback loop. Our observations do not directly address this question but suggest the utility of future studies defining the responding T-cells with respect to both antigen specificity and proviral HIV-1 DNA. Our efforts to define the influence of the timing of the HZ rash on the CSF HIV-1 RNA concentration and its relation to that of plasma did not provide a clear definition of this factor, though the numbers were small and the timing of LPs inconsistent. However, the aggregate cross-sectional experience with escape and discordance together with the longitudinal observations show that the effect of HZ on CSF HIV-1 RNA, in parallel with the inflammatory reaction, is protracted, lasting up to the 5 month limit of our study definition and beyond one year in the longitudinal example in Figure 4. The elevation of the age-corrected CSF NfL concentrations to above normal or baseline limits in more than half of all the HZ cases, including out to 150 days in one of the individuals with CSF escape, also documents the prolonged time course of HZ, and the longitudinal case shows how this injury also can parallel the CSF escape. We did not detect a clear overall correlation of either the NfL concentration or the distance of the HZ rash from the LP needle with the magnitude of the effect on CSF HIV-1 RNA. While we presume that the inflammatory focus, including the focus of HIV-1 RNA release, is anatomically circumscribed, our data were insufficient to define this quantitatively and may have been obscured by ready diffusion of HIV-1 RNA and other analytes within the CSF space.

The two non-HZ escape cases included in this analysis exhibited features that align with the four individuals with HZ-related secondary escape, but also included some differences. Most notably, they shared the association of the CSF HIV-1 RNA concentrations with the WBC counts, enough to fit into the linear correlation between these two key variables (Figure 3). However, the HSV-2 encephalitis case differed in the background CD4+ T-cell count (130 cells per µL), a low level suggesting that this unusual infection might be classified as ‘opportunistic’. However, despite this exceptional CD4+ T cell count, this individual exhibited a robust CSF pleocytosis in common with the other escape cases, consistent with the role of inflammation in augmenting the CSF HIV-1 RNA level. The individual with neuroborreliosis had 320 blood CD4+ T-cells per µL, closer to the range of the four HZ escape cases. Further virological characterization of these two non-HZ cases along with a broader range of non-opportunistic and opportunistic CNS infections needs to be done to define any differences compared to escape in the HZ setting. Similarly, further study of the CSF in the untreated discordant individuals and relationship of CSF viral populations to those of plasma in this setting (where, unlike in the escape cases, concurrent plasma virus concentrations are sufficient for RNA studies) as well as comparing these to non-discordant and to other untreated individuals should also enhance the studies of secondary escape alone.

HZ provides a relatively common and useful model of secondary escape. Even in the presence of suppressive ART its incidence is higher in PLWH than in the general population, providing an opportunity to extend these retrospective observations. Secondary escape in HZ at least partially resembles asymptomatic escape and raises the question of whether HZ itself, with or without rash [68], or other mild CNS infections or inflammatory conditions might play a role in some episodes of asymptomatic escape, particularly where escape is sustained over several weeks or months [38].

### Limitations

Our study had several limitations. The number of individuals in each of the four HZ groups was relatively small, particularly in the two treated groups. It was also a retrospective study of a single cohort, though this particular cohort study had the good fortune to include protocol LPs and CSF analyses of HIV-1 RNA, and therefore afforded a valuable opportunity to examine this issue. The timing of LPs after HZ was not set by *a priori* design to carefully define the frequency or time course of the effect of HZ on local HIV-1 infection, but rather was determined mainly by the protocol-specified schedule and non-protocol clinical decisions. This almost random timing in relation to the HZ episode had the advantage of showing the prolonged period of pleocytosis and CSF escape in some subjects but did not allow consistent assessment of the time course of this escape as would a study specifically focused on this issue. The longitudinal observations before and after HZ in some individuals were useful in providing interesting background to the impact of ART and its disruption by HZ, but only afforded a preliminary view of the time course of HZ effects. The two non-HZ cases appear to support the findings in HZ but may involve some different mechanisms despite their seeming similarity, so their integration into more unifying conclusions needs to be cautious. In the untreated individuals the zoster episode led to initiation of ART, and consequently follow-up and natural history without HIV-1 treatment was not available nor was follow-up information on CSF after treatment (for example, whether CSF HIV-1 RNA remained elevated after the resolution of plasma viremia as might be predicted if discordance effects were the same as those of escape). On the other hand, these untreated individuals exhibiting parallel relations of CSF WBCs to CSF HIV-1 RNA provide an important and provocative perspective on secondary escape related to their likely shared mechanisms.

## Conclusions

This study defines secondary CSF escape as a coherent entity and HZ as a relatively common and useful model of it. The study also shows some of the parallel features of CSF escape and discordance in treated and untreated individuals, respectively, that likely indicate similar pathogenetic mechanisms, particularly the interaction of reactive inflammation and CSF HIV-1 RNA levels. The results suggest that the robust local or regional inflammatory response to VZV gives rise to secondary HIV-1 production that is reflected in CSF. They also demonstrate that secondary escape is an entity worthy of further study, including virological and immunological dissection that may contribute to a broader understanding of some of the features of a contributing source of CSF HIV-1 infection.

## Acknowledgements

The authors would like to acknowledge the medical staff at the Sahlgrenska University Hospital, Gothenburg Sweden.

## Conflicts of Interest

LH, RWP, DF, MG none. HZ has served at scientific advisory boards for Roche Diagnostics, Wave, Samumed and CogRx, has given lectures in symposia sponsored by Biogen and Alzecure, and is a co-founder of Brain Biomarker Solutions in Gothenburg AB, a GU Ventures-based platform company at the University of Gothenburg (all outside submitted work).

## Notes

**Funding:** This study was funded by grants from the Swedish state under the agreement between the Swedish government and the county councils, the ALF-agreement (ALFGBG-717531) and National Institutes of Health/National Institute of Neurological Diseases and Stroke grant R01NS094067. HZ is a Wallenberg Scholar, additionally supported by grants from the Swedish Research Council (2018-02532), the European Research Council (681712) and the Swedish state under the agreement between the Swedish government and the county councils, the ALF-agreement (ALFGBG-720931).

